# A multiverse analysis of stomach-brain coupling in humans

**DOI:** 10.64898/2026.04.22.720086

**Authors:** Trinh Thi Thuy Ngo, Tzu-Yu Hsu (徐慈妤), Niall W. Duncan

**Affiliations:** Graduate Institute of Mind, Brain and Consciousness, Taipei Medical University, Taipei, Taiwan; Institute of Cognitive Neuroscience, National Central University, Taoyuan, Taiwan

**Keywords:** Interoception, Stomach, Brain, Brain-body, Methodology, Reproducibility

## Abstract

The gastric-brain axis is a burgeoning field of neuroscience; however, inferences from neuroimaging research are often constrained by the high dimensionality of methodological choices potentially leading to disparate outcomes. This study addresses such concerns by performing a multiverse analysis of gastric-brain coupling in humans. We systematically evaluated 5,184 unique analytic pipelines using electroencephalography (EEG) and electrogastrography (EGG) data to survey the methodological sensitivity of gastric-brain coupling. Our results reveal that whilst analytic decisions influence the magnitude of observed coupling, at the group level the phenomenon remains relatively consistent across the parameter space. High inter-individual variance can, however, be observed. Coupling was observed in the alpha, theta, and beta bands, with the latter two bands showing coupling at the largest number of electrodes. Coupling across frequency bands was primarily seen in medial electrodes, with some left lateral coupling also observed. Overall, these findings suggest that EEG-EGG based measurement of gastric-brain coupling is likely to be consistent across a significant range of plausible analytic pipelines at the group-level in healthy participants, providing useful context for future studies.

## Introduction

The connection between the digestive system and the brain is a topic of increasing interest. A burgeoning literature is elucidating how these organ systems interact and how these interactions can influence our experience and behaviour [1,2]. Levels of investigation range from circulating factors arising from gut biota to the effects of bidirectional direct innervation between the systems. Importantly, evidence indicates that interactions between the systems may play a role in pathological conditions [3,4]. There are thus both scientific and clinical motivations for developing our understanding of this axis as part of a wider research programme probing the relationship between the brain and the body [1,5,6].

In this context, neuroimaging studies in humans have begun to map patterns of dynamic influences of stomach contractile activity on the brain [7]. This gastric-brain coupling was first shown with the amplitude of brain alpha oscillations being linked to the slow rhythm of the stomach [8]. Subsequently, the spatial extent of gastric-brain coupling was mapped with functional MRI (fMRI), showing a network of brain regions that are coupled to the contractile rhythm of the stomach [9,10]. A similar spatial pattern of gastric-brain coupling has also been seen with MEG measures of brain activity [11]. These studies together introduce a complex interaction between the stomach and brain and open up important avenues of future research.

Although such neuroimaging studies of the gut-brain axis are likely to play an important role in our understanding of how the brain and body interact, there are known issues within the imaging field related to analytic flexibility. Across different modalities, it has been highlighted that the high dimensionality of possible methodological choices available when analysing neuroimaging data may lead to variable analytic outcomes [12,13]. Empirical work has shown that such variability can result in false negatives, false positives, or outcomes in the opposite direction from the true effect [14–20]. This can lead to a lack of robustness in a research domain and, particularly if connected to publication bias, may lead to invalid inferences being made by the research community.

With gastric-brain coupling research at a relatively early stage, we therefore sought to quantify the ‘multiverse’ of analytic outcomes [21,22]. By systematically varying preprocessing and estimation parameters, we aimed to identify which nodes of the analysis pipeline are most influential in determining the presence or strength of observed coupling. It is hoped that surveying the influence of analytic choices on outcomes in this way can contribute to a strong methodological foundation that future research can build upon for robust and replicable results. This may be of particular importance for the gastric-brain field as the need to combine multiple complex measures from two separate organ systems presents a large set of possible decisions to researchers.

For this work we chose to focus on EEG-based measures of brain activity for estimating the gastric-brain coupling. This was for three reasons. Firstly, EEG appears to be a fast growing modality for gastric-brain coupling, with a number of studies using that modality being published around the time of writing [23–27]. Secondly, EEG is a convenient approach for studies of pathological states. Thirdly, EEG is also more accessible to those living in lower-resourced areas, which may help reduce potential bias in our understanding of the gastric-brain system from selective sampling [28,29]. It should be noted, however, that the use of EEG has downsides in the context of measuring potential gastric-brain coupling. In particular, the low spatial resolution limits separation between different brain networks, which may show coupling within specific frequency bands [11]. This could limit the effects that can be detected.

Simultaneous EEG and electrogastrography (EGG) data were therefore acquired from a group of healthy young adults. These data were then entered into a multiverse analysis and the resulting set of outcomes further analysed to establish the methodological sensitivity of any group-level evidence for coupling between the EEG and EGG data.

## Methods

### Participants

Forty-six adults were recruited for the study. Participants were required to have: no history of eating disorders; no acute or chronic gastrointestinal disorders; no current or past diagnosis of psychiatric or neurological disorders; normal or corrected-to-normal vision; and a body mass index (BMI) of 18-25. Of the 46 recruited, six were excluded due to a lack of a reliable EGG signal. This left a sample of 40 participants (20 female; mean age = 27.7 ± 3.2 years; mean BMI = 21.0 ± 1.8). The study was approved by the Taipei Medical University Institutional Review Board (N202408044). Participants provided written informed consent and were financially compensated for their participation.

### Experimental procedure

Participants were asked to refrain from eating in the four hours prior to their session, and to limit fluid intake in the two hours prior. They were also asked to not consume nicotine on the day of the study or alcohol from the day before. Upon arriving, participants were informed about the study and consent was obtained. After this, EEG and EGG electrodes were placed on the head and abdomen. Participants were seated in a semi-recumbent position that allowed them to fully relax their abdominal muscles to facilitate the measurement of the gastric slow-wave signal. Data acquisition was conducted in a lit room. All sessions began at 16:00 to minimise variation due to circadian or metabolic factors.

During the study, each participant completed two resting state sessions lasting ten minutes each. During these periods, they were instructed to keep their eyes open, relax, and minimize movement. A fixation cross was located in their eyeline, upon which they were asked to maintain their gaze. A short break was provided between the sessions. The experimental procedures, including preparation, questionnaires, EEG, and EGG acquisition, required approximately 1.5–2 hours.

### EGG acquisition and data preparation

Gastric myoelectrical signals were recorded using a nine-channel OpenBCI Cyton biosensing board (OpenBCI, New York City, USA). Adhesive Ag/AgCl electrodes were positioned following a standardised abdominal montage for capturing gastric slow-wave potentials [30]. In more detail, the first electrode (E1) was placed approximately 2 cm above the umbilicus on the midline, with two additional electrodes (E2 and E3) positioned superiorly along the same axis at intervals corresponding to one-third and two-thirds of the distance to the xiphoid process. Another vertical column of electrodes was placed beneath the midpoint of the left clavicle. These electrodes (E6 and E7) were aligned horizontally with the E1 and E2. When necessary, the upper left electrode (E7) was shifted medially to avoid interference from the costal margin. Two intermediate electrodes (E4 and E5) were placed midway between these vertical columns, positioned at the midpoints between E1-E2 and E2-E3, respectively. A reference electrode was located contralateral to the midline at a site symmetric to E5, and a ground electrode was placed on the lower left abdomen above the iliac crest. Details of abdominal electrode placement are given in Figure 1A. Data were sampled at 250 Hz.

**Figure 1:**
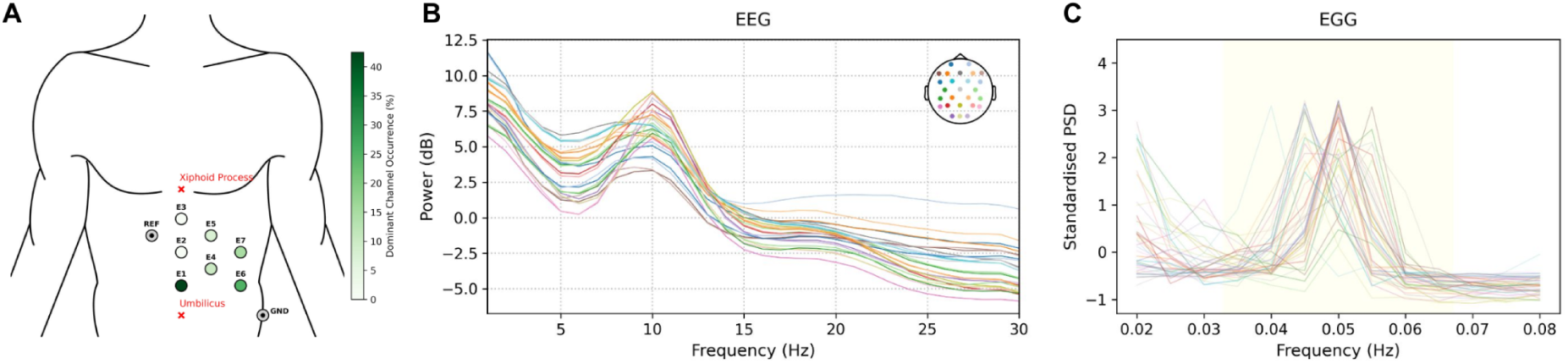
(A) Illustration of the location of the seven abdominal electrodes (E1–E7) used for electrogastrography (EGG) recordings. Marker colour shade reflects the proportion of participants for whom that electrode is identified exhibiting the highest peak power within the gastric slow wave range (0.033–0.067 Hz). (B) Participant power spectra for included EEG data. (C) Participant power spectra for included EGG data. As raw signal amplitude naturally varies across individuals [30], spectra were calculated from standardised time EGG time series to allow comparison of peaks across participants.

Prior to analysis, EGG recordings were cropped to the trigger indicating the start of the recording session. Slow drifts were removed from each channel using a polynomial regression (Legendre basis functions, orders 1–3). After removing faulty cycles, participants had a median of 29 (range = 24 - 32) valid gastric cycles included in the analysis (see Table S1 and Figure S2 for details).

### EEG acquisition and data preparation

Brain electrophysiological signals were acquired using a 32-channel Brain Products system configured according to the international 10–20 electrode placement scheme. Four electrooculography (EOG) electrodes were positioned around the eyes to facilitate removal of ocular artifacts. Two electrodes were placed on the mastoid processes as reference channels. Data were sampled at 1000 Hz and electrode impedances were kept below 10 kΩ for all participants. Continuous data were recorded using Brain Vision Recorder software (Brain Products, Munich, Germany).

Prior to data analysis, EEG signals were downsampled to 250 Hz to match the EGG. BrainVision files were imported into MNE-Python (v1.7) [31] and cropped to the trigger indicating the start of the recording session (see Supplementary Methods for details). A 1-30 Hz FIR band-pass filter was applied using MNE’s zero-phase forward–backward filtering, selecting a linear-phase design to avoid phase distortion in subsequent phase–amplitude coupling (PAC) analyses. Bad channels were detected via visual inspection and replaced using spherical spline interpolation.

### Multiverse analysis pipelines - Overview

Analysis of the data was split into three stages: EGG preprocessing, EEG preprocessing, and coupling quantification. Each of these stages then has multiple decision points. For the multiverse analysis, sets of plausible decisions for each decision point were determined based upon approaches previously reported in relevant literature. Note that limited sets of plausible decisions were used rather than aiming for an exhaustive potential set as the limited set will allow a demonstration of whether outcomes are sensitive to methodological decisions or not while remaining computationally viable. An overview of decision sets is given in Table 1. These pipelines were run for a set of four EEG frequency bands: delta (1-4 Hz), theta (4-8 Hz), alpha (8-13 Hz), and beta (13-30 Hz). In total, 5,184 different analysis pipelines were calculated. Pipelines were implemented in Python (v3.11) through a combination of MNE-Python (v1.7), SciPy (v1.15.3), NumPy (v2.3.5), and NiTime (v0.9).

**Table 1:** Overview of multiverse decision steps. ICA = independent component analysis; FIR = finite impulse response; STD = standard deviation; IIR = infinite impulse response; IQR = interquartile range; MAD = median absolute deviation; MI = Modulation Index; CLC = circular-linear correlation.

| Decision point |  | Options (N) | Decision options | Specifics |
| --- | --- | --- | --- | --- |
| EGG | PSD estimation | 2 | Welch method,<br>Multi-taper method | Gastric frequency range:<br>0.033–0.067 Hz |
|  | Channel selection | 2 | Max-power channel | Channel with maximum power |
| | | | Average of “good” channels | Average of channels with $\geq 90\%$ of max power |
| | Filter | 2 | FIR, IIR | Bandwidth $\pm 0.015$ Hz around peak frequency |
| | Recording rejection criteria | 1 | Normogastric proportion check | Recordings with $< 70\%$ of cycles in 2–4 cpm range were excluded |
|  | Cycle rejection method | 3 | STD + phase monotonicity check |  |
|  |  |  | MAD + phase monotonicity check |  |
|  |  |  | IQR + phase monotonicity check |  |
|  | Threshold (k) | 3 | 2.5; 3.0; 3.5 | Threshold level for duration-based outlier rejection in each cycle rejection method |
| **EGG combinations** |  | **72** |  |  |
| EEG | EEG band | 4 | Delta, Theta, Alpha, Beta |  |
|  | Referencing strategy | 2 | Common average reference, linked mastoid (A1–A2) |  |
|  | ICA method | 2 | Manual (visual inspection), Automated* (ICLabel) | Automated approach only applied to average reference due to ICLabel requirement |
| **EEG combinations** |  | **12** |  |  |
|  | Surrogate computation | 3 | One-block shuffle, five-block shuffle, circular time shift |  |
|  | Coupling computation | 2 | MI; CLC |  |
| **PAC combinations** |  | **6** |  |  |

One run out of the two acquired was selected for the multiverse analysis based upon which had the better data quality. This was established based upon the following criteria: the presence of a clear alpha peak in the EEG data following basic preprocessing; a clear peak within the gastric slow wave range (i.e., 0.033–0.067 Hz) in the EGG; and more than 70% of EGG cycles falling within the normative human normo-gastric range (15–30 s) [30]. Two further quality assurance steps were applied to the EGG data of the selected runs. The first of these involved quantifying the proportion of time within the run where the normogastric rhythm was dominant (compared to bradygastric and tachygastric rhythms). A threshold of 70% of the run was applied, based upon prior literature [32]. Following this, the standard deviation of participant’s gastric cycle durations was calculated. Any participant with a cycle duration standard deviation of greater than six was excluded. One subject was found to have a dominant normogastric rhythm in below 70% of the run and was excluded (Figure S3). All 39 of the remaining participants had a cycle duration SD below six (Figure S4). Group EEG and EGG power spectra for the selected runs are shown in Figure 1B&C.

### Multiverse analysis pipelines - EGG

For the processing of the EGG data, the EGG power spectrum was calculated within the gastric rhythm range of 0.033–0.067 Hz [30] using either (i) Welch’s method (window length = 200 s, 75% overlap) or (ii) a multi-taper estimator implemented in the NiTime python package. Selection of the dominant channel(s) was then done by either (i) selecting the channel with the maximal spectral power, or (ii) according to an averaged-channel approach that took the mean signal across channels whose peak power exceeded 90% of the maximum. The selected channel(s) were then filtered using either (i) a finite impulse response (FIR) filter with a passband of ± 0.015 Hz centered on the individual normo-gastric peak frequency or (ii) a fourth-order Butterworth band-pass filter centered on the individual normo-gastric peak. Instantaneous phase and amplitudes were then obtained using the analytic signal of the Hilbert transform.

Gastric cycles were identified at negative phase-wrap transitions (π → −π), and their durations were computed. Artifact detection was implemented as a multiverse of nine cycle-based rejection configurations (3 statistical criteria x 3 thresholds). Central tendency and dispersion were estimated using one of three methods: standard deviation (SD), median absolute deviation (MAD), or interquartile range (IQR). For each method, admissible durations were defined as lying within k ∈ {2.5, 3.0, 3.5} units of the center. Cycles were rejected if their durations exceeded method-specific bounds or if their phase progression was non-monotonic (e.g., noise-induced phase reversals). Only cycles satisfying both criteria were retained. A binary valid mask was generated for each pipeline, marking timepoints belonging to accepted cycles. This mask was later applied to the EEG, such that EEG signal amplitude was extracted only during intervals in which gastric activity was deemed physiologically plausible.

### Multiverse analysis pipelines - EEG

Following the basic EEG data preparation steps outlined above, the first decision steps for the EEG pipelines was a combination of which reference to use (mastoid or the common average) and the method for ICA artifact removal. For data using the common average reference, both (i) a manual identification of artifact components based on visual inspection of topographies and component time courses, or (ii) automatic classification using ICLabel, implemented via *MNE-ICALabel* [33,34]. Components classified as *eye, muscle, line noise, heart,* or *channel noise* were removed, while those labeled as *brain* or *other* were retained. Only manual identification was used for data using the mastoid reference as ICLabel requires an average reference. Band-limited EEG signals were then obtained for the relevant canonical frequency bands [11,35] via finite impulse response (FIR) filtering. The amplitude envelope was extracted using the Hilbert transform.

### Multiverse analysis pipelines - Coupling

The coupling between the phase of the EGG and the amplitude of the EEG was calculated through two different approaches: (i) the modulation index (MI) [8,36], and (ii) circular-linear correlation (CLC) [11]. MI quantifies coupling by assessing how far the empirical distribution of amplitudes across phase bins deviates from uniformity. The gastric phase time series was divided into 18 equidistant bins spanning the interval [-*π*, *π*], following previous relevant literature [8,25]. For each bin, the mean EEG amplitude was computed, yielding a phase–amplitude distribution. The MI is then defined as the Kullback–Leibler divergence between the observed distribution and a uniform distribution, normalised by the maximum possible divergence [36]. The CLC coefficient was calculated based on correlations between EEG amplitude and the sine and cosine components of EGG phase across the same 18 bins [37].

Three surrogate methods were used to generate null distributions of PAC values: one-block swapping, circular time shifting, and five-block shuffling. For the one-block method, the EEG amplitude time series was cut at a random time point and the two resulting segments were concatenated in reverse order. For circular time shifting, the amplitude time series was shifted by a random temporal offset and wrapped around at the recording boundaries. For five-block shuffling, the amplitude time series was divided into five contiguous segments that were randomly reordered. These manipulations preserve the amplitude distribution and spectral properties of the EEG signal while disrupting its temporal alignment with the gastric phase. One-thousand surrogate coupling values were calculated for each pipeline. Empirical PAC values were then standardized relative to its corresponding surrogate distribution as

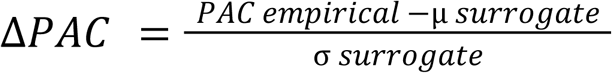

where *μ*_surrogate_ and *σ*_surrogate_ denote the mean and standard deviation of the surrogate PAC distribution, respectively. Positive ΔPAC values indicate stronger empirical coupling than expected by chance.

### Specification curve analysis

To evaluate whether gut–brain PAC was present across the set of preprocessing pipelines, we conducted a specification curve analysis (SCA) following the general framework proposed by Simonsohn et al. [21]. The input to the SCA consisted of subject-level channel-wise ΔPAC values obtained for every preprocessing pipeline. Group-level coupling across participants was then tested through a one-sided Wilcoxon signed-rank test of ΔPAC being greater than zero.

Three inferential summary statistics were computed from the set of Wilcoxon outcomes across all pipelines [21]. Firstly, we calculated the median Wilcoxon effect size *r* across all specifications for a given channel, indexing the typical magnitude of coupling across the multiverse. Secondly, we calculated the proportion of specifications yielding statistically significant positive coupling, indexing the consistency with which that channel exceeded chance-level PAC across pipelines. Thirdly, we computed a Stouffer-type summary statistic by converting specification-level *p*-values to Z scores and averaging them across pipelines, thereby capturing the aggregate strength of evidence for that channel across the full specification space. Statistical significance of these channel-level SCA summary statistics was evaluated using permutation-based null distributions. Under the null hypothesis of no systematic coupling, subject-level ΔPAC values were sign-flipped within the channel, while the same subject-specific sign was preserved across all pipelines for that channel within a permutation iteration. For each permutation, the full set of channel × pipeline Wilcoxon tests was recomputed, followed by recalculation of the three SCA summary statistics. Empirical *p*-values were then obtained by comparing the observed statistics against their corresponding permutation-derived null distributions.

### Processing decision influence

A linear mixed-effects model (LMM) was used to investigate which preprocessing decisions systematically influenced gut–brain coupling strength. Taking ΔPAC values across all pipelines as the dependent variable, the different decision steps and frequency bands were modeled as fixed effects. Participant ID and EEG channel were included in the model as random effects. The model was specified as:

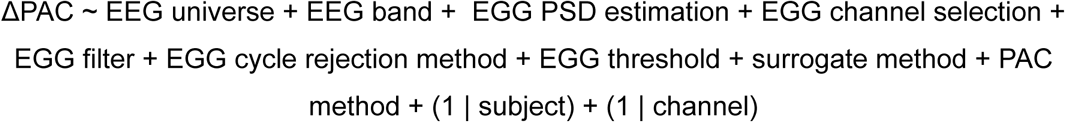

The analysis was conducted in R (v4.5.1) with the *lme4* package. Follow-up comparisons were performed using estimated marginal means implemented in the *emmeans* package. Pairwise contrasts were corrected using the Holm method to control the family-wise error rate. Owing to the large sample size, post-hoc contrasts were evaluated using asymptotic z-tests. Whereas the SCA assessed the robustness of gut–brain coupling across the full multiverse of preprocessing pipelines, the LMM provided a complementary analysis aimed at identifying which analytic choices were systematically associated with stronger or weaker PAC estimates.

## Results

### Overall EEG-EGG coupling

Evidence for consistent EEG-EGG coupling was found when considering all channels and frequency bands (**Figure 2**). CP1 showed the most consistent evidence, showing an elevated proportion of significant pipelines (19.6%, p = 0.009) and mean Stouffer’s Z (0.55, p = .039), although its median correlation was not significant (r = .087, p = 0.055). Two additional channels survived one test each: F4 for mean Stouffer’s Z (0.56, p = 0.038) and FC5 for the proportion of significant pipelines (14.7%, p = 0.023). Overall, significant findings were sparse and were distributed across frontal and centroparietal/parietal electrodes, with the strongest evidence observed at CP1.

**Figure 2:**
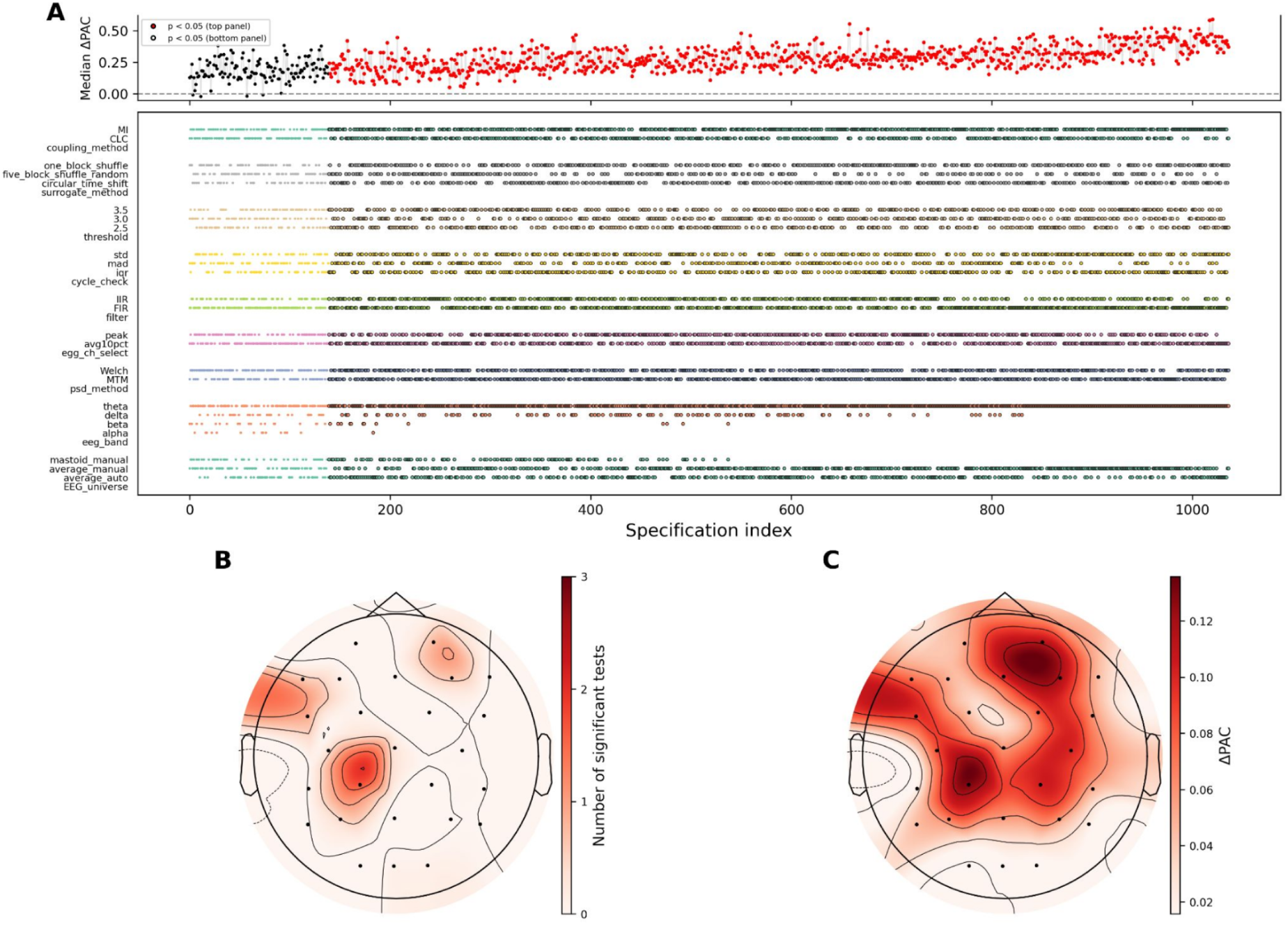
Outcome from multiverse analysis across all frequency bands. A) Specification curve illustrating the top 20% of pipelines averaged across channels with robust coupling. B) Scalp topography of the number of significant inferential tests passed by each EEG channel C) Topography of average coupling strength across pipelines and frequency bands.

The LMM analysis of influences on overall ΔPAC identified a number of factors that were significantly related to outcomes (see Table 2 for model coefficients). For EGG processing, PSD method, channel-selection method, and cycle-rejection criterion all showed differential ΔPAC across analytic choices. Post-hoc tests showed that ΔPAC was higher when the selected EGG channel was estimated by averaging EGG channels exceeding 90% of peak power compared to selecting a single dominant channel, while the multi-taper method resulted in a higher ΔPAC than Welch’s approach. For cycle handling, the standard deviation criterion resulted in higher ΔPAC than alternative methods.

**Table 2:** Linear mixed-effects model on channel-level ΔPAC from all analytical pipelines. Reference levels were common average reference with automated ICA classification, Multi-taper PSD estimation method, IIR filtering, max-power channel selection, IQR cycle-rejection criterion, threshold value of 3.5, theta band, one-block swapping surrogate method, and MI coupling method. Coefficients represent differences relative to these reference levels. Positive coefficients indicate higher ΔPAC than the reference category. ICA = independent component analysis; FIR = finite impulse response; IIR = infinite impulse response; IQR = interquartile range; MAD = median absolute deviation; MI = Modulation Index; CLC = circular-linear correlation. SD = standard deviation; SE = standard error. CIs are 95% Wald confidence intervals.

| Fixed effects | Estimate | SE | 95% CI | *t* | *p* |
| --- | --- | --- | --- | --- | --- |
| Intercept | 0.129 | 0.044 | [0.04, 0.218] | 2.93 | .006 |
| Average reference + manual ICA | -0.005 | 0.001 | [-0.008, -0.003] | -4.47 | < .001 |
| Linked mastoid reference + manual ICA | -0.02 | 0.001 | [-0.023, -0.018] | -17.3 | < .001 |
| Welch's method | -0.006 | 0.001 | [-0.008, -0.004] | -6.57 | < .001 |
| FIR filtering | -0.001 | 0.001 | [-0.003, 0.001] | -1.30 | .194 |
| EGG Averaged-channel strategy | 0.018 | 0.001 | [0.016, 0.020] | 18.7 | < .001 |
| MAD | -0.011 | 0.001 | [-0.013, -0.009] | -9.21 | < .001 |
| SD | 0.005 | 0.001 | [0.002, 0.007] | 3.79 | < .001 |
| Threshold = 2.5 | -0.001 | 0.001 | [-0.003, 0.002] | -0.55 | .585 |
| Threshold = 3.0 | -0.003 | 0.001 | [-0.005, 0.000] | -2.13 | .033 |
| Alpha band | -0.033 | 0.001 | [-0.036, -0.031] | -24.2 | < .001 |
| Beta band | -0.023 | 0.001 | [-0.026, -0.021] | -16.8 | < .001 |
| Delta band | -0.111 | 0.001 | [-0.114, -0.108] | -80.78 | < .001 |
| Surrogate method - Circular time shift | -0.001 | 0.001 | [-0.003, 0.002] | -0.35 | .724 |
| Surrogate method - Five-block shuffle | -0.04 | 0.001 | [-0.04, -0.038] | -33.5 | < .001 |
| Coupling method - CLC | -0.001 | 0.001 | [-0.003, -0.001] | -1.43 | .152 |
| Random effects | Estimate | SD |  |  |  |
| Participant variance (intercept) | 0.074 | 0.27 |  |  |  |
| Channel variance (intercept) | 0.001 | 0.04 |  |  |  |
| Residual variance | 1.24 | 1.11 |  |  |  |

For EEG processing, the reference and denoising approach was found to have an influence on ΔPAC, with a common average reference with automated ICA being associated with highest values.

For surrogate generation, the one-block swapping and circular time-shift methods did not differ (p = 0.724), whereas both produced higher ΔPAC than the five-block shuffle method (both Holm-adjusted p < 0.001). All post hoc comparisons are reported in Table 3

**Table 3:**
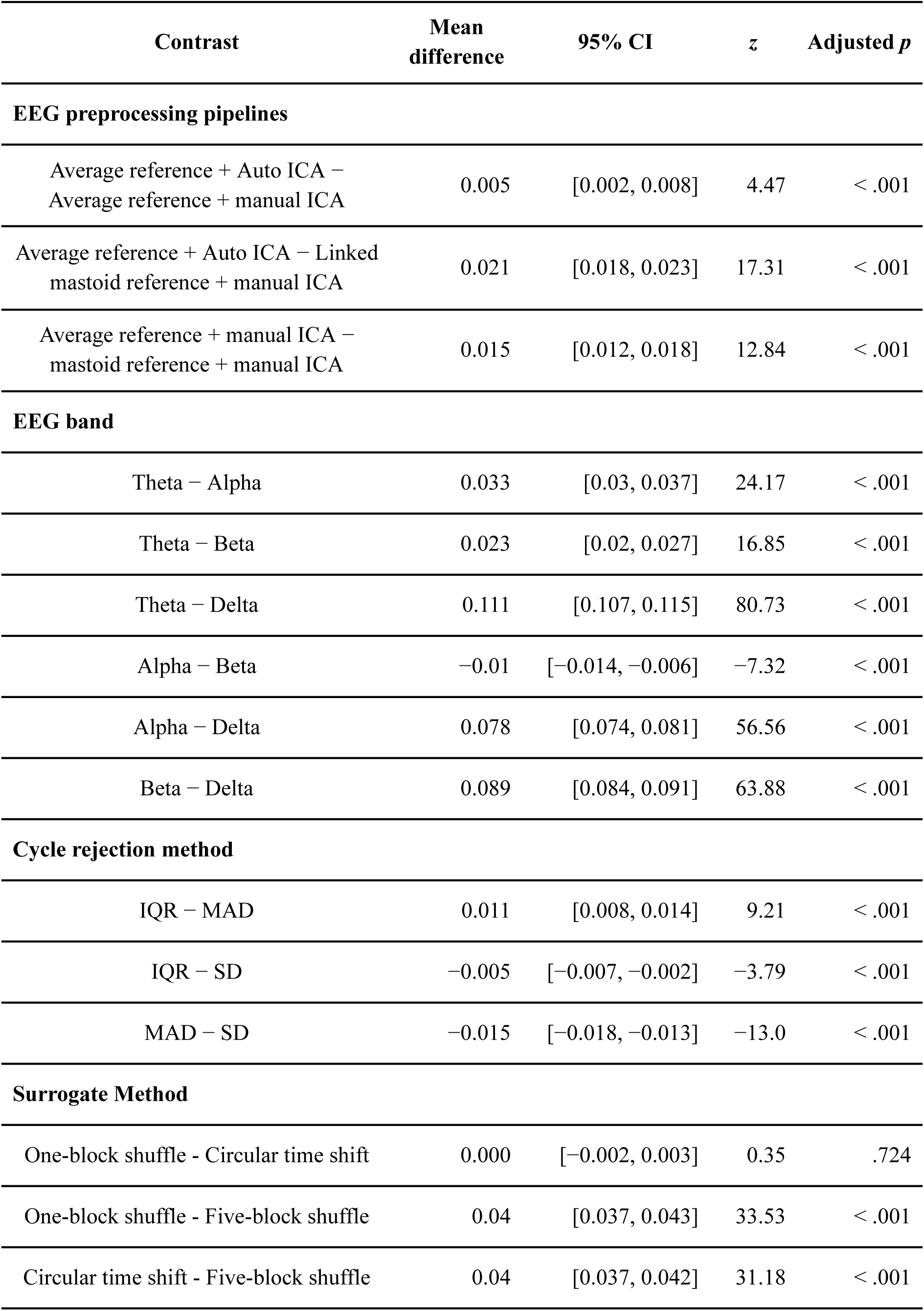
Pairwise comparisons of estimated marginal means for significant multiverse analysis factors.

We note, though, that fixed analytic factors accounted for only 0.2% of the variance in ΔPAC (marginal *R*² = 0.002), whereas the full mixed model explained 5.9% (conditional *R*² = 0.059).

It may also be noted that the analysis outcome is strongly influenced by data quality, as shown by a supplementary comparison between the selected and rejected runs. An LMM analysis showed strongly reduced ΔPAC values in the rejected run (*β* = −0.117, *t* = −167.43, *p* < 0.001), meaning that no significant coupling is found in those runs assessed to have less prominent alpha peaks and EGG activity within the normogastric range (see Figure S5).

### Band specific EEG-EGG coupling

The LMM analysis also identified a difference in ΔPAC values across frequency bands. Post hoc analyses revealed that ΔPAC differed across all EEG frequency bands (all Holm-adjusted p < 0.001), with theta producing the highest values, followed by beta, alpha, and delta. This pattern is also seen when running SCA on the bands separately, where we see no channels with significant coupling across pipelines in the delta and alpha range (Figure 3 & see Figures S6-S9 for band-specific SCA). Theta-band coupling exhibited a predominantly left frontocentral–centroparietal region, with consistent evidence at FC5, C3, and CP1 (all surviving the three inferential tests). Beta-band coupling was observed at right frontoparietal electrodes, with CP2, Fz, and P4 surviving all three inferential tests, F4 surviving two inferential tests, and additional evidence at FC2 and P3 from one inferential test.

**Figure 3:**
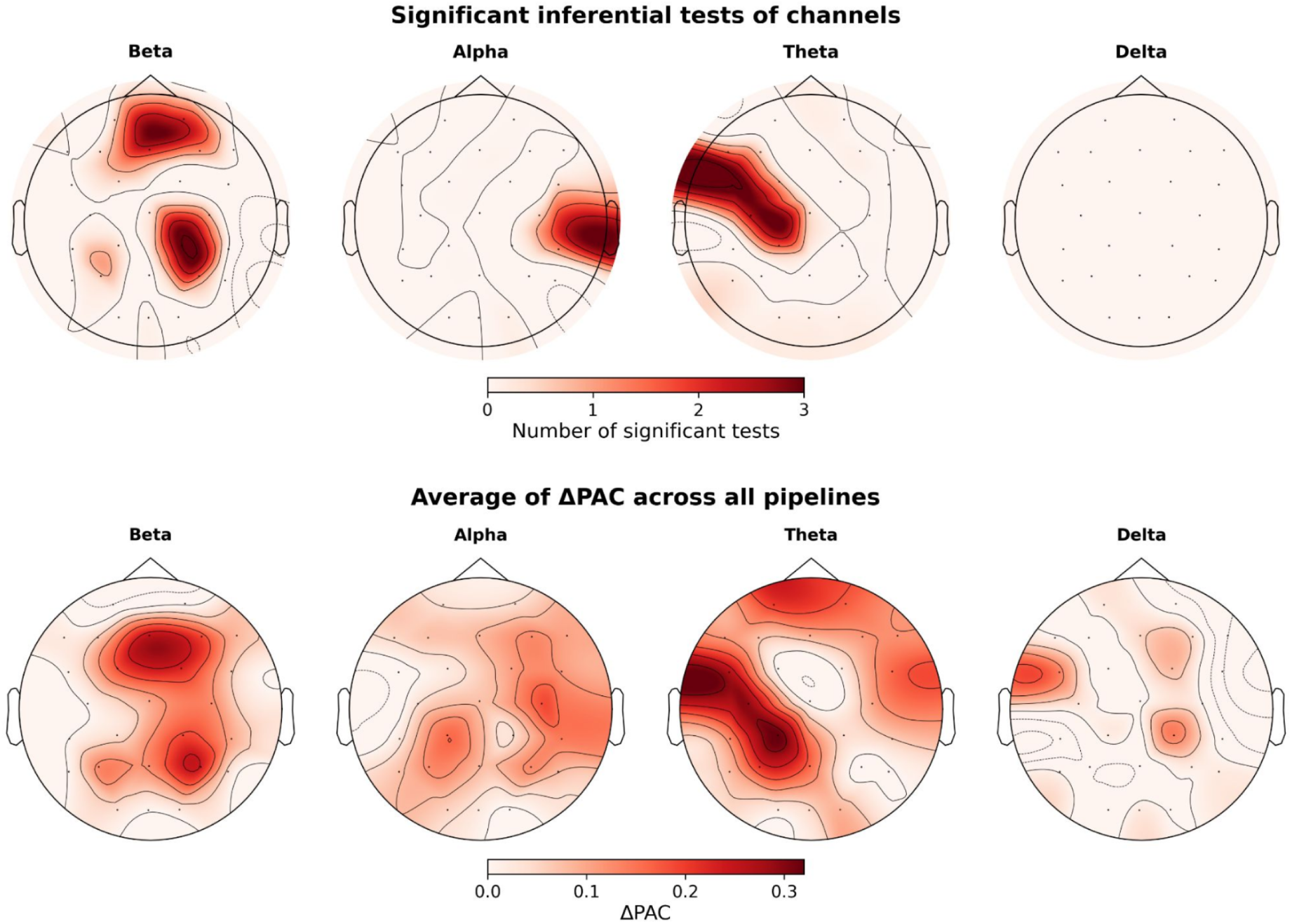
Band-specific multiverse outcomes for delta, theta, alpha, and beta bands. Upper panels show scalp topography of the number of significant inferential tests passed by each EEG channel. Lower panels show topography of average coupling strength across pipelines.

## Discussion

The present study employed a multiverse analysis to systematically evaluate the methodological sensitivity of gastric-brain coupling in humans, as measured through EEG and EGG. By traversing 5,184 unique analytic pipelines, we aimed to determine the extent to which reported coupling is a product of specific methodological choices or represents an analytically stable phenomenon. Our results demonstrate that while analytic decisions do influence the magnitude of observed coupling, evidence for EEG-EGG coupling can be observed across a significant subset of the multiverse of analytical choices studied. This coupling is, however, relatively sparse and weak in this dataset, with large inter-individual variation.

Our identification of coupling in all frequency bands other than delta fits with other work showing coupling across a wide frequency range [11]. We do see some differences from prior results, however, where delta coupling has been reported [11] and where alpha coupling has been more centred [8]. These prior results were obtained with MEG and so the contrast may reflect differences in the capabilities of that methodology compared to EEG. In particular, the lack of spatial specificity inherent to EEG means that anatomically localised frequency-specific coupling may be obscured. Alternatively, our work used canonical frequency bands for the analysis, meaning that more frequency specific coupling may have been missed. A more extensive multiverse analysis varying this factor may therefore be justified.

Topographically, the channels showing the strongest evidence for coupling were distributed across frontal and centroparietal/parietal electrodes, with CP1 emerging as the most consistent measurement location across frequency bands. Although scalp electrodes cannot be interpreted as direct indicators of underlying cortical sources, this distribution would appear to be broadly compatible with previous MEG and fMRI studies describing a distributed “gastric network” involving sensorimotor and higher-order cortical regions [9,11,38]. This potential consistency with prior work may be further supported by the frequency specific topographies of significant coupling, where coupling in the beta and theta bands follow a general frontal and central distribution.

Beyond investigating how sensitive coupling outcomes are to analytical choices, our results may provide some guidance for optimising future EEG-EGG research pipelines. The LMM analysis indicated that certain choices systematically yielded higher coupling estimates. For EGG processing, we found that estimating power spectra using the multi-taper method and utilising an averaged-channel approach (taking the mean of channels exceeding 90% of peak power) were superior to Welch’s estimation or selecting a single dominant channel.

This average-channel approach differs from that suggested in prior work [30]. Regarding EEG, the combination of a common average reference and automated ICA denoising was associated with the highest coupling values. It can be noted that the second best approach was an average reference with manual rejection, pointing to the average reference being generally preferable. The method for generating null distributions for measuring coupling was also found to be relevant, with one-block swapping and circular time-shift surrogates generating similar outcomes and both exceeding the five-block shuffle approach. This fits well with recommendations that surrogates for continuous recordings should minimize distortion of phase and amplitude dynamics, with single-point block swapping preferred over resampling at multiple cut points [39]. Importantly, it must be noted that although differences between approaches were found, the ultimate magnitude of these differences was generally not high.

Large inter-individual variation in gastric-brain coupling was seen in this dataset. Notably, the regression model suggests that this variation had a considerably larger influence on ΔPAC values than did methodological choices. Prior work with MEG has estimated that only around 8% of the variance in alpha amplitude was accounted for by the gastric oscillation [8]. This relatively small effect may be diluted further when using EEG and so apparent individual variation may arise from a lack of measurement precision [40]. Alternatively, the individual variability may reflect true differences in communication between the stomach and brain between participants. Although efforts were made to ensure consistency in the data recording context for all participants, there are still factors which could influence the level of input to the brain and how that affects ongoing brain activity. Importantly, gastric signals do not reach the cortex directly. Instead, gastric information is conveyed to the central nervous system through vagal and spinal afferent pathways [7]. Vagal afferents terminate primarily in the nucleus tractus solitarius, from which visceral information can influence hypothalamic and distributed brain networks [41]. Thus, variability in autonomic state or in the sensitivity and efficacy of these afferent pathways could influence how strongly gastric rhythms modulate cortical activity. One potential source of such variability is differences in circadian time between participants. Circadian rhythms are endogenously generated but are entrained by multiple environmental and behavioral cues, including light exposure, habitual sleep timing, feeding schedules, and physical activity [42–44]. The central circadian system coordinates peripheral physiology through sympathetic and parasympathetic autonomic pathways [45–47], together with hormonal, thermal, and behavioral signals [48].

Consequently, participants recorded at the same clock time, as was done here, may nevertheless have differed in internal circadian phase and autonomic state. Overall, we can note that the presence of identifiable gastric slow waves and cortical oscillations should be regarded as a prerequisite for assessing coupling but are in themselves not evidence that strong coupling must occur.

The use of EEG in this study was motivated by its increasing prevalence in the field and its availability in clinical contexts. Our demonstration that gastric-brain coupling could be observed within this dataset across a meaningful subset of analytic choices points towards the potential of EEG as a tool for probing brain-body interactions in diverse settings.

Importantly, as EEG is more accessible than fMRI or MEG, it allows for the inclusion of participants from lower-resourced areas, thereby reducing potential selective sampling bias in our understanding of the gastric-brain system [28]. The relative consistency across analytic pipelines that we observed suggests that EEG-based gastric-brain coupling may have the potential to serve as a useful tool for investigating gastrointestinal or neurological disorders characterised by dysregulated brain-body communication. However, as noted above, use of EEG limits the spatial specificity of the measurement of brain activity. Some evidence suggests that different brain networks may be associated with gastric activity in frequency-specific ways [11]. The mixing of signals between networks that could occur due to low spatial resolution would therefore potentially obfuscate such specificity. This means that the results obtained may only reflect a subset of gastric-brain associations and may omit others that exist. Any results should therefore be interpreted with this context in mind and method choice should follow the specific aims of a given investigation.

### Limitations

Several limitations of the current study should be acknowledged. Firstly, our participant sample was restricted to individuals with a BMI between 18 and 25 and no history of gastrointestinal or psychiatric disorders. While this was necessary for establishing a baseline in a healthy population, it is unclear if the same level of analytic robustness would be found in clinical populations where signals might be more attenuated or noisy. Secondly, a proportion of our initial sample (approximately 13%) was excluded due to poor EGG signal quality. This exclusion may bias results towards particular subsets of the population from whom clear signals can be obtained. Thirdly, our work presents outcomes from multiple SCA analysis across the 26 EEG channels. This raises questions about the appropriate approach to potential corrections for multiple comparisons for this analysis type. It must be noted that the results given here have not had such a correction applied. Fourthly, 1000 surrogates were used in the analyses. It is possible that the choice of surrogate number could influence outcomes. However, it was not computationally plausible for us to extend our multiverse to include this factor. This leaves the effect of this decision point open. Finally, this work focused on resting-state data; future multiverse analyses could investigate if analytic flexibility has a greater influence during active tasks or following metabolic challenges (e.g., meal ingestion).

## Conclusion

In conclusion, this multiverse analysis helps add to the methodological foundation for the study of the gastric-brain axis. Gastric-brain coupling could be observed at the group-level in this dataset across a meaningful set of analytic decisions. This coupling was, however, relatively weak, with large inter-individual variability. The epistemological issue motivating multiverse analyses also remains. We as investigators do not know whether a reported outcome from any particular analysis pipeline is a chance product of the specific decisions made or not [22,49]. As such, a more general adoption of this form of analysis may be justified within the brain-body research field to support more robust inferences.

## Acknowledgments

Funding for this research was provided by grants from Taiwan’s National Science and Technology Council to NWD (113-2423-H-038-002-MY3; 110-2628-H-038-001-MY4) and TYH (113-2410-H-008-082, 114-2410-H-008-066, 115-2410-H-008-051-MY3).

## Author contributions

TTNT: Conceptualization, Formal analysis, Investigation, Methodology, Visualization, Writing – original draft

TYH: Methodology, Supervision, Writing – original draft

NWD: Conceptualization, Methodology, Supervision, Writing – original draft

## Data and code availability

Code and data supporting this work are available at https://doi.org/10.17605/OSF.IO/8T9MZ.

## Competing interests

The authors declare no competing interests.

A multiverse analysis of stomach-brain coupling in humans

## Supplementary methods

### EEG-EGG recording synchronisation

To achieve temporal correspondence between the cortical and gastric recordings, a hardware-based synchronisation system was implemented (Figure S2). The synchronisation mechanism is centered on an isolated trigger interface designed for use with the OpenBCI Cyton biosensing board (Electools, China). This trigger board incorporates optical isolation circuitry and digital input–output channels, enabling safe bidirectional communication with external acquisition systems while electrically isolating participant-connected hardware. The trigger board was directly mounted onto the OpenBCI 9-channel biosensing unit and linked to the EEG amplifier through a DB25-to-terminal block connected to the EEG amplifier USB 2 port (BUA). This configuration provided a stable TTL-level pathway between the two independent acquisition systems. Optical isolation components (rated to approximately 4000 V) ensured full galvanic separation, eliminating the risk of current transfer between devices. A dedicated digital channel (D17) on the trigger board was used as the synchronization marker. This line is hardwired to a physical push-button. Activating the button generated a TTL pulse that was dispatched simultaneously along two routes:

1. To the OpenBCI system, where it was recorded as a digital event embedded directly in the EGG data stream; and
2. To the BrainAmp amplifier, where it was received by the trigger input of the BrainVision system and registered at the corresponding sampling time in the EEG recording.

During preprocessing, EEG and EGG signals were cropped at the first detected D17 pulse, yielding precisely matched temporal windows for subsequent phase–amplitude coupling analysis.

## Supplementary figures

**Figure S1:**
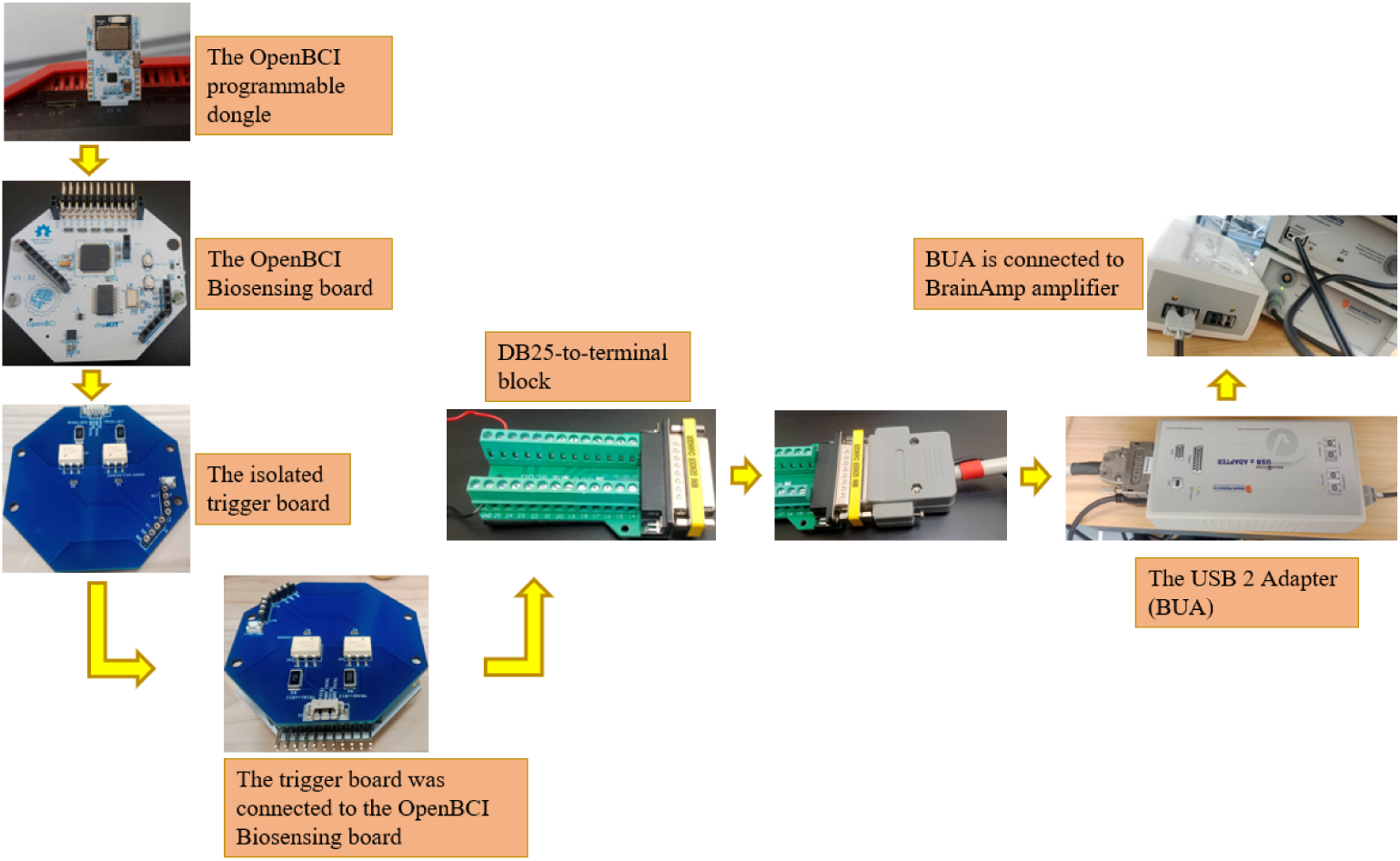
Schematic diagram of the hardware-based synchronization setup for simultaneous EEG and EGG acquisition

**Figure S2:**
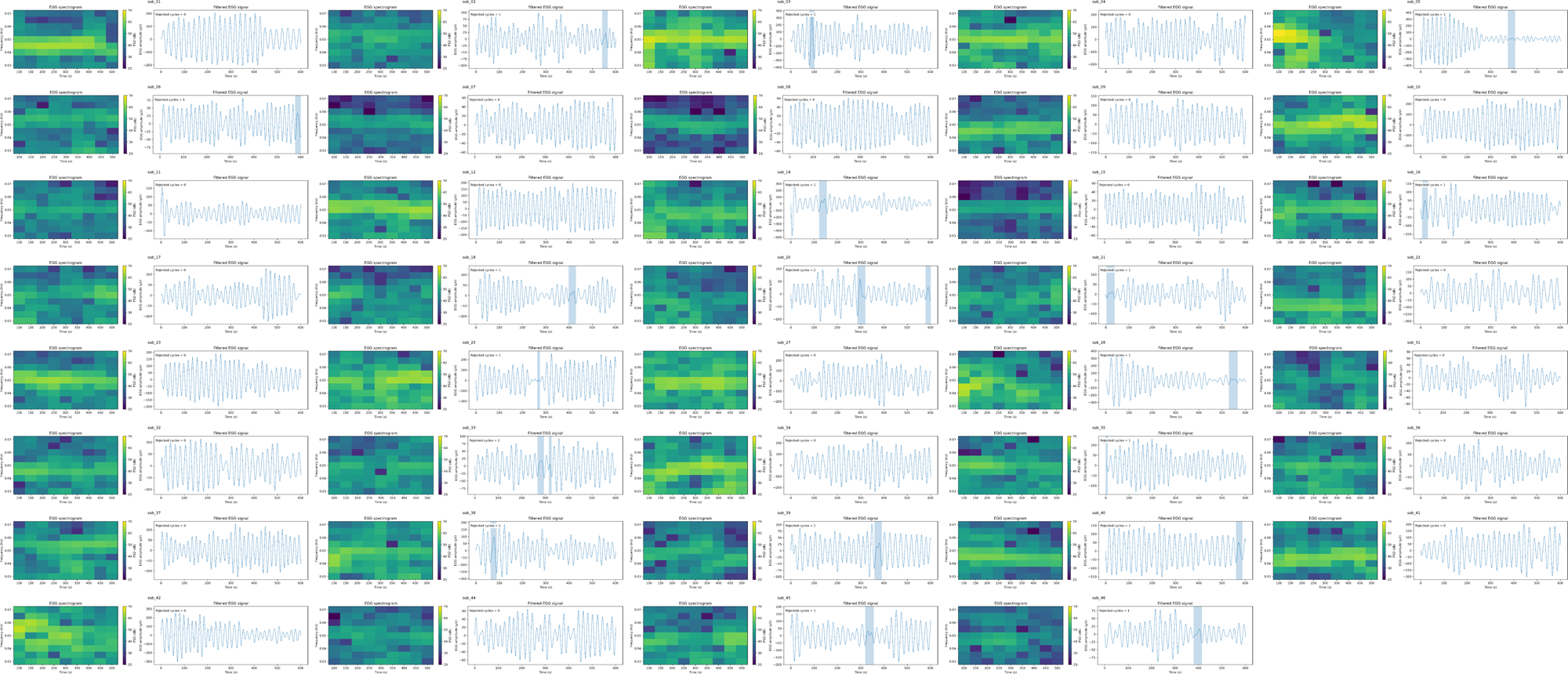
EGG time series for included runs for all participants. Excluded cycles are indicated by the blue shaded areas.

**Figure S3:**
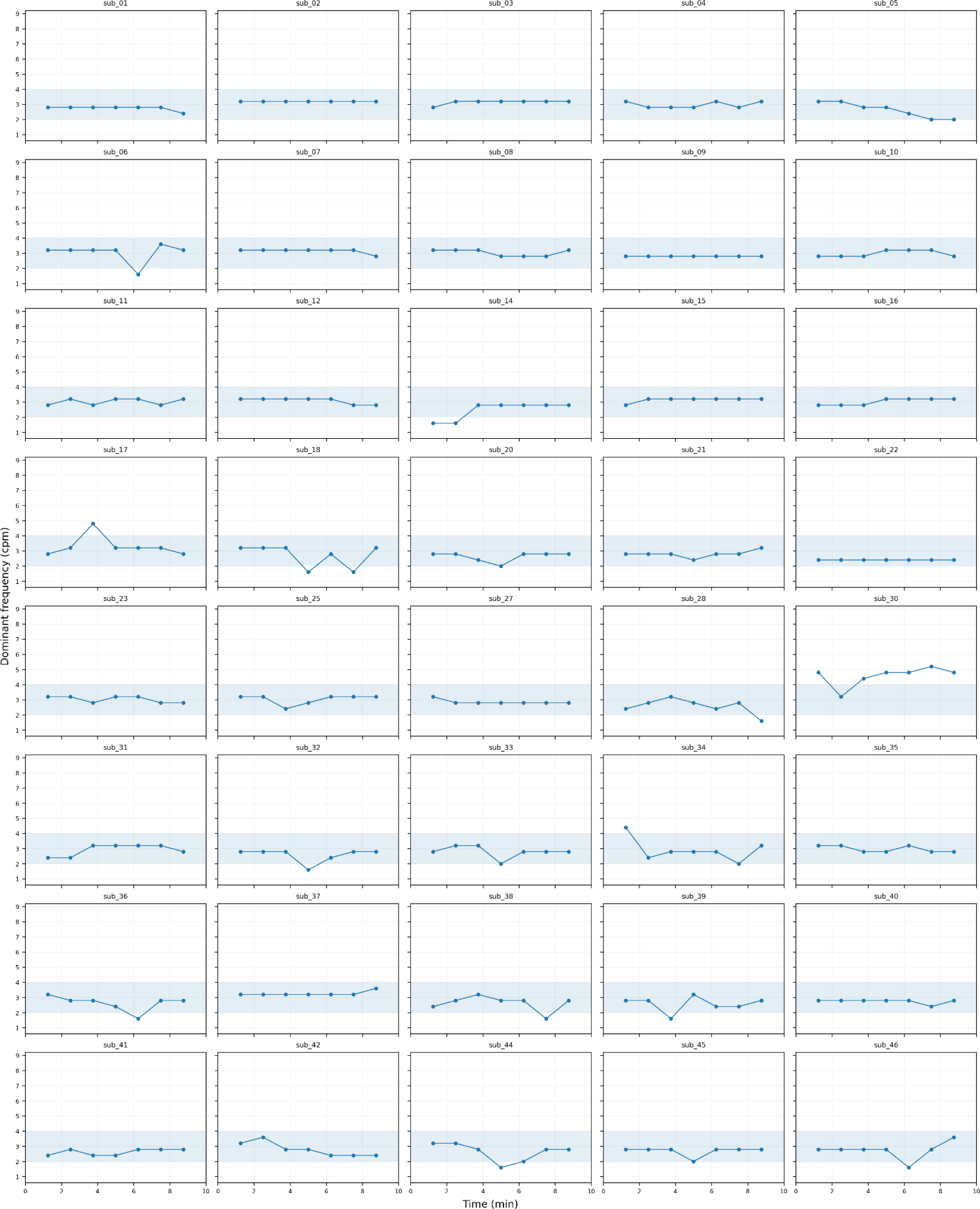
Temporal dynamics of electrogastrographic dominant frequency across all participants. One participant (sub_30) was excluded from the subsequent analyses because fewer than 70% of the analysis windows exhibited a DF within the normogastric range.

**Figure S4:**
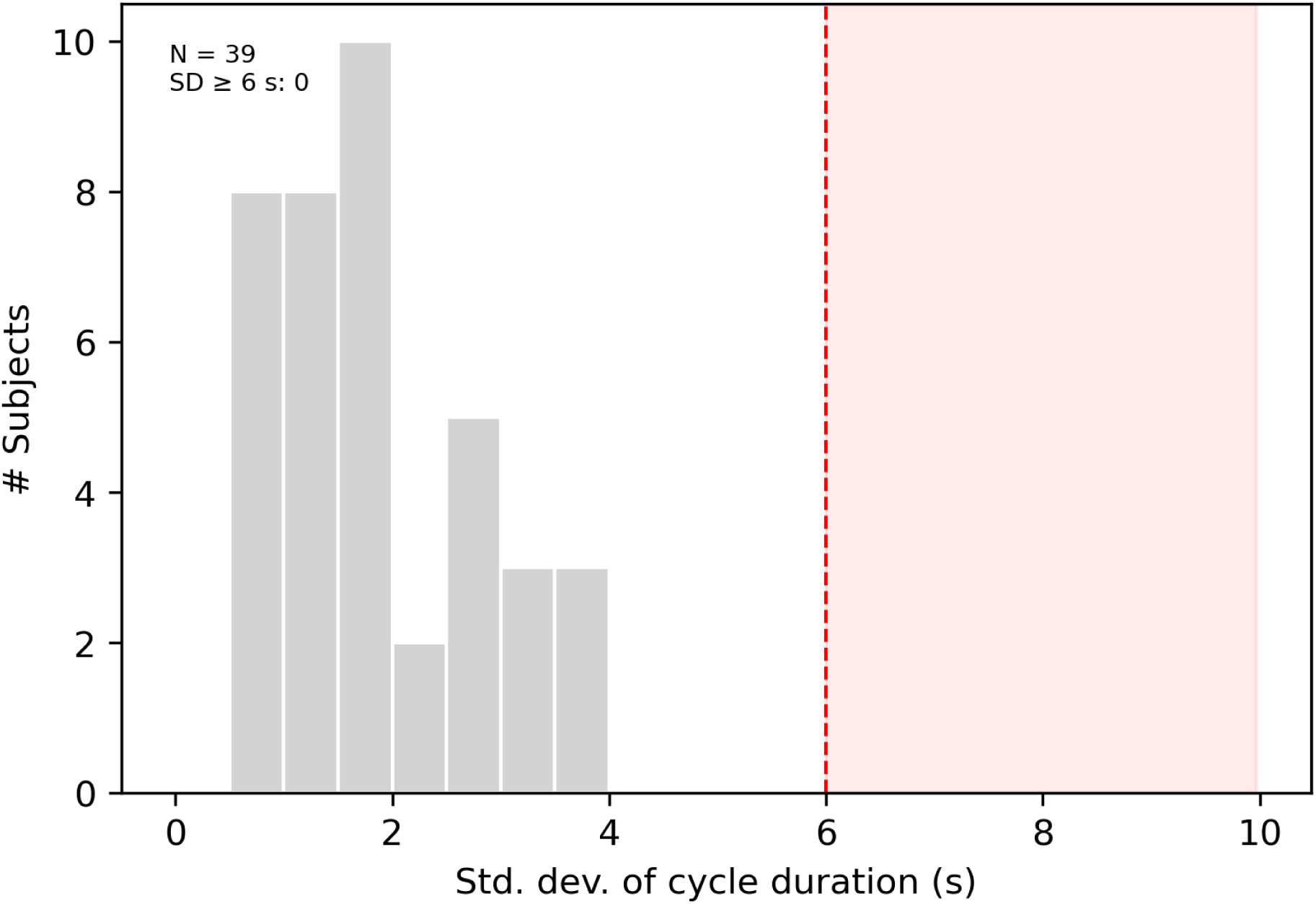
The distribution of the SD of gastric cycle duration among 39 participants retained after quality control.

**Figure S5:**
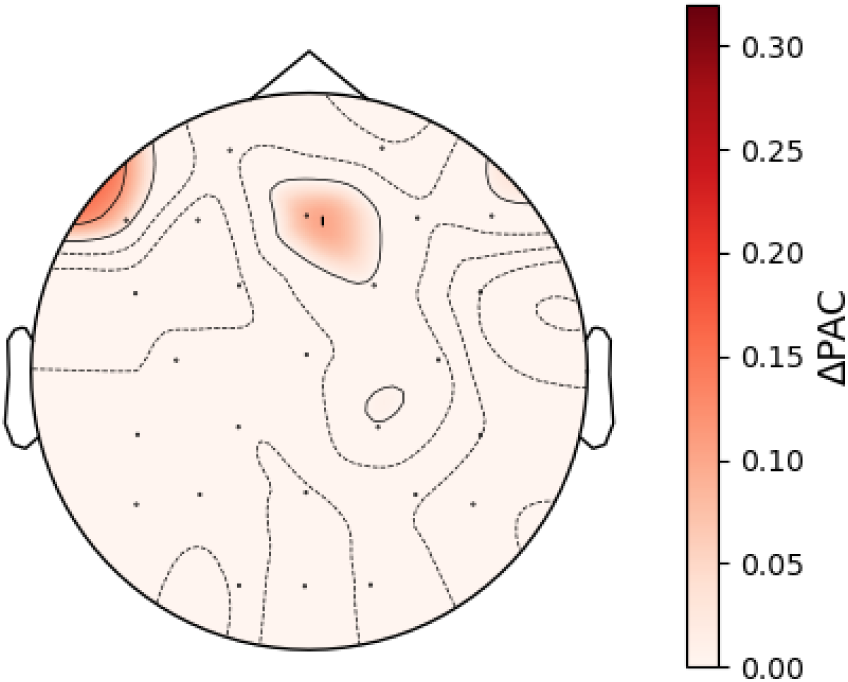
Topographical plot showing average coupling strength across pipelines and frequency bands for the runs not selected for the main analysis. No electrodes have coupling that passes any inferential tests.

**Figure S6:**
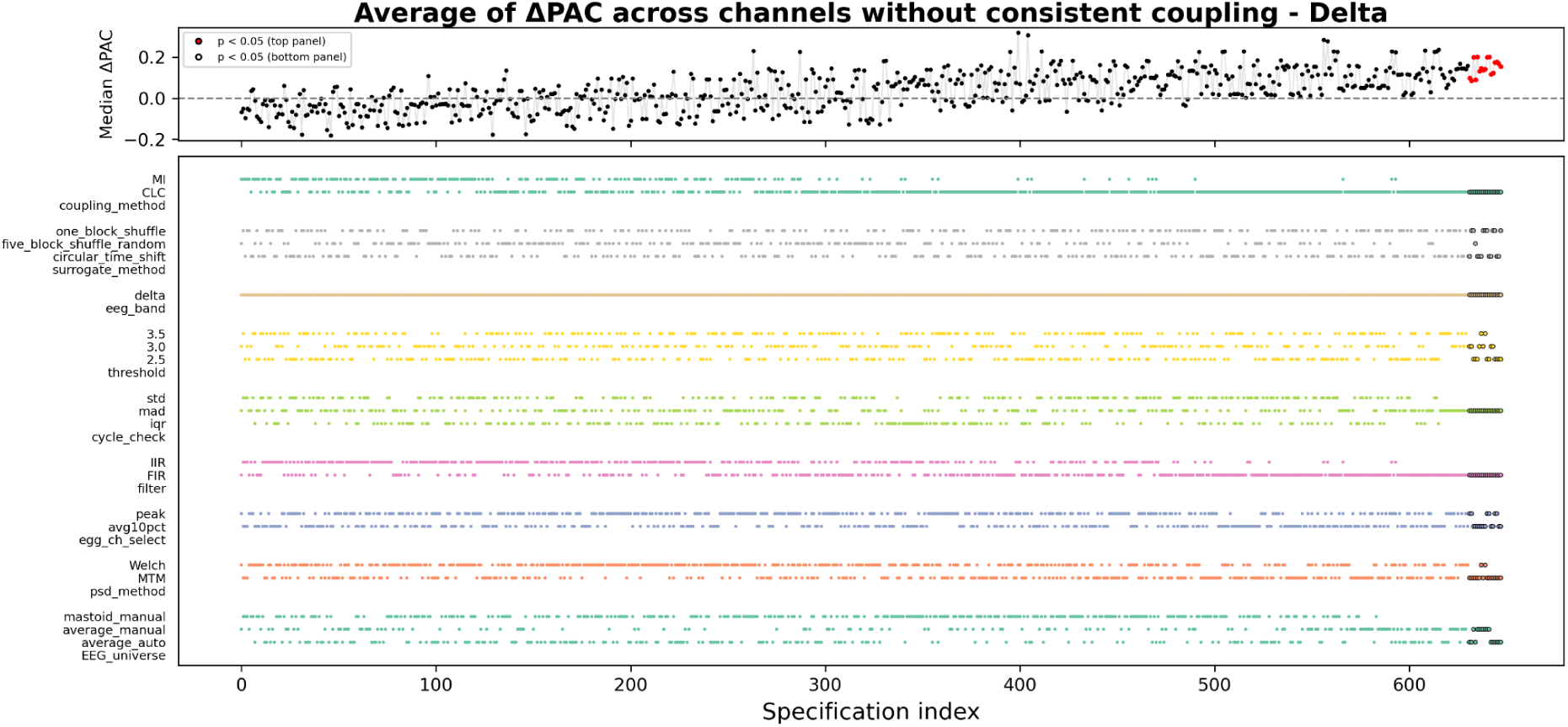
Specification curve illustrating the top 20% of pipelines averaged across channels without robust coupling for the delta band specific multiverse analysis.

**Figure S7:**
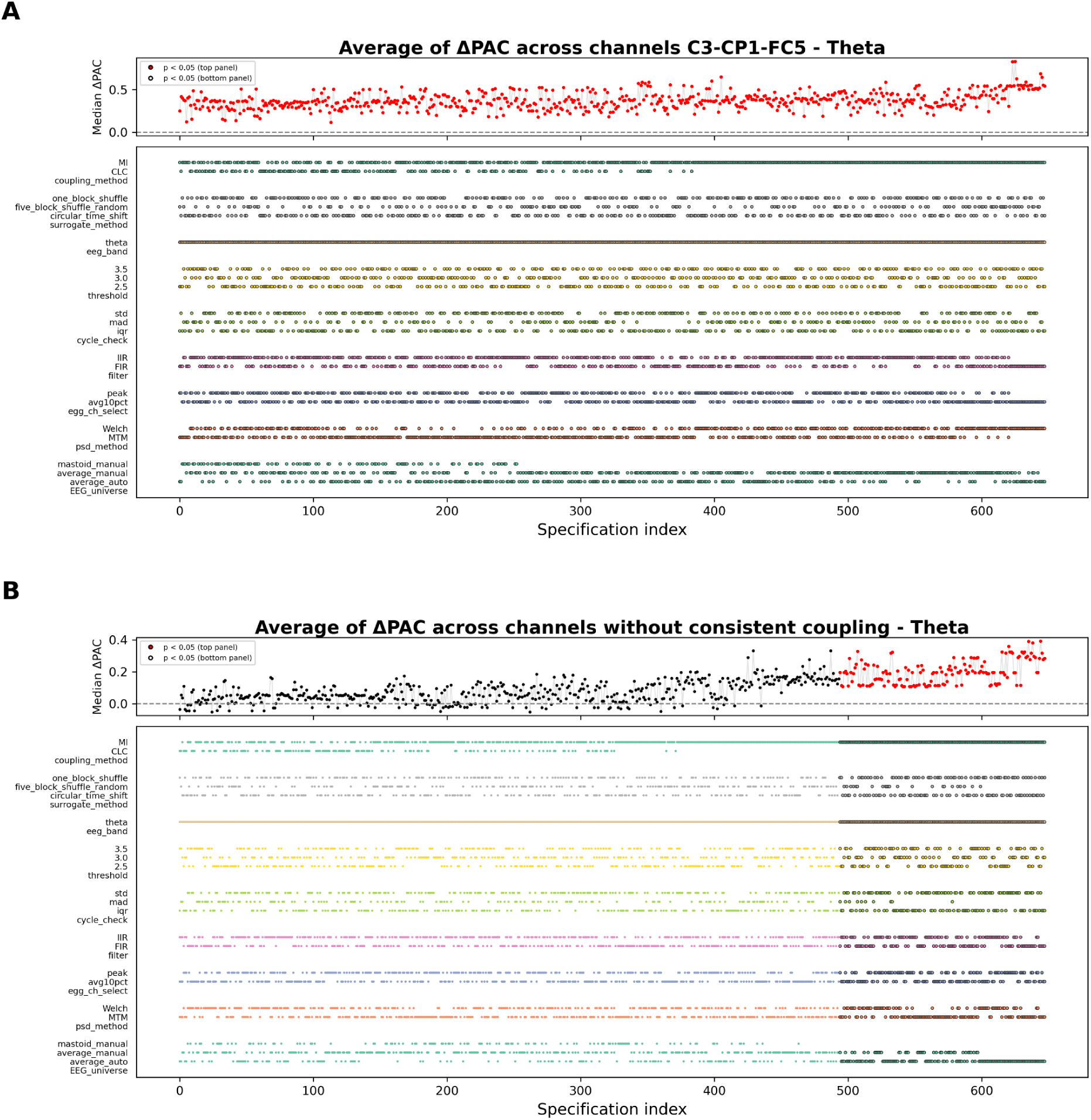
Specification curve illustrating the top 20% of pipelines averaged across A) channels with robust coupling and B) without robust coupling for the theta band specific multiverse analysis.

**Figure S8:**
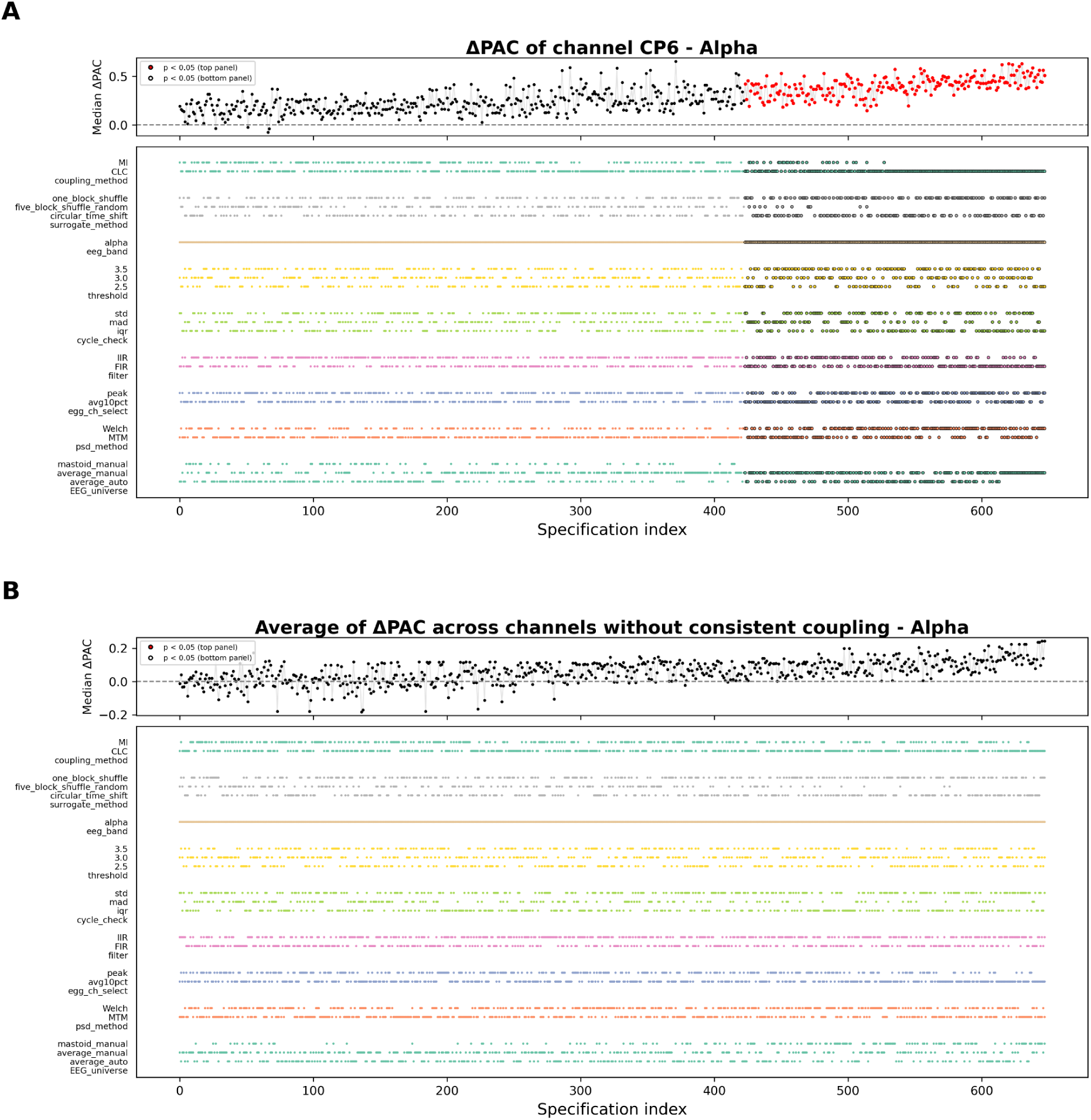
Specification curve illustrating the top 20% of pipelines averaged across A) channels with robust coupling and B) without robust coupling for the alpha band specific multiverse analysis.

**Figure S9:**
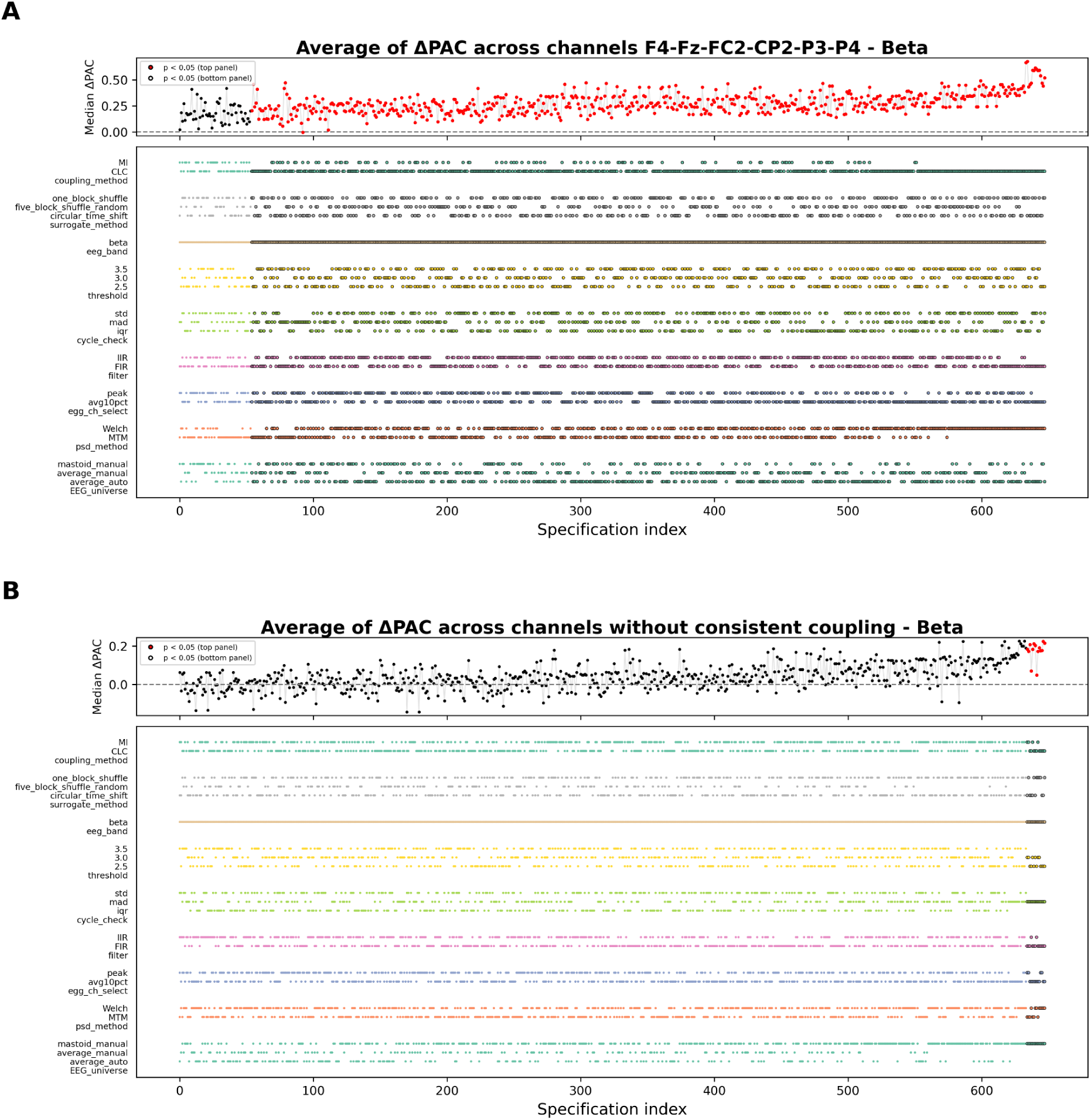
Specification curve illustrating the top 20% of pipelines averaged across A) channels with robust coupling and B) without robust coupling for the beta band specific multiverse analysis.

## Supplementary tables

**Table S1:** Details of EGG time series for the runs included in the analysis.

| Subject id | Total EGG cycles | Rejected EGG cycles | Remaining time (s) |
| --- | --- | --- | --- |
| sub-01 | 25 | 0 | 579.2 |
| sub-02 | 32 | 1 | 572.9 |
| sub-03 | 30 | 1 | 579.7 |
| sub-04 | 29 | 0 | 587.0 |
| sub-05 | 29 | 1 | 567.1 |
| sub-06 | 32 | 0 | 595.9 |
| sub-07 | 30 | 0 | 581.2 |
| sub-08 | 29 | 0 | 586.7 |
| sub-09 | 27 | 0 | 590.1 |
| sub-10 | 29 | 0 | 593.9 |
| sub-11 | 29 | 0 | 585.9 |
| sub-12 | 30 | 0 | 596.1 |
| sub-14 | 27 | 0 | 596.4 |
| sub-15 | 30 | 0 | 588.0 |
| sub-16 | 28 | 1 | 561.3 |
| sub-17 | 29 | 0 | 572.2 |
| sub-18 | 29 | 1 | 565.6 |
| sub-20 | 25 | 1 | 553.9 |
| sub-21 | 28 | 0 | 595.8 |
| sub-22 | 24 | 0 | 588.7 |
| sub-23 | 29 | 0 | 592.3 |
| sub-25 | 30 | 2 | 575.9 |
| sub-27 | 28 | 0 | 587.2 |
| sub-28 | 24 | 1 | 549.5 |
| sub-31 | 29 | 1 | 560.3 |
| sub-32 | 26 | 0 | 589.0 |
| sub-33 | 28 | 1 | 567.0 |
| sub-34 | 27 | 0 | 591.8 |
| sub-35 | 29 | 0 | 591.8 |
| sub-36 | 29 | 0 | 597.4 |
| sub-37 | 31 | 0 | 584.9 |
| sub-38 | 28 | 0 | 580.4 |
| sub-39 | 27 | 1 | 554.8 |
| sub-40 | 26 | 0 | 595.9 |
| sub-41 | 26 | 0 | 594.5 |
| sub-42 | 31 | 0 | 585.0 |
| sub-44 | 28 | 2 | 534.4 |
| sub-45 | 27 | 0 | 585.4 |
| sub-46 | 26 | 1 | 556.2 |

## Notes

https://doi.org/10.17605/OSF.IO/8T9MZ

### Competing Interest Statement

The authors have declared no competing interest.

### Summary of Updates

Additional data quality checks were added. More analytical factors were included in the multiverse analysis.

